# Catalytically Enhanced Cas9 through Directed Protein Evolution

**DOI:** 10.1101/2020.07.01.183194

**Authors:** Travis H. Hand, Mitchell O. Roth, Chardasia L. Smith, Emily Shiel, Kyle N. Klein, David M. Gilbert, Hong Li

## Abstract

The Clustered Regularly Interspaced Short Palindromic Repeat (CRISPR)-Cas9 system has found widespread applications in genome manipulations due to its simplicity and effectiveness. Significant efforts in enzyme engineering have been made to improve the CRISPR-Cas9 systems beyond their natural power with additional functionalities such as DNA modification, transcriptional regulation, and high target selectivity^1–10^. Relatively less attention, however, has been paid to improving the catalytic efficiency of CRISPR-Cas9. Increased catalytic efficiency may be desired in applications where the currently available CRISPR-Cas9 tools are either ineffective^4, 11–14^ or of low efficiency such as with type II-C Cas9^15–18^ or in non-mammals^19, 20^. We describe a directed protein evolution method that enables selection of catalytically enhanced CRISPR-Cas9 variants (CECas9). We demonstrate the effectiveness of this method with a previously characterized Type IIC Cas9 from *Acidothermus cellulolyticus* (AceCas9) with up to 4-fold improvement of in vitro catalytic efficiency, as well as the widely used *Streptococcus pyogenes* Cas9 (SpyCas9), which showed a 2-fold increase in homology directed repair (HDR)-based gene insertion in human colon cancer cells.

---

Functional CRISPR-Cas9 enzymes are composed of a Cas9 protein, a CRISPR RNA (crRNA) and a trans-activating crRNA (tracrRNA). The crRNA and tracrRNA can often be replaced by a chimeric single-guide RNA (sgRNA)^21^.The sgRNA or crRNA contains a complementary region that base pairs with the target DNA (protospacer). Cas9 possesses multiple domains: a nucleic acid recognition domain (REC), a PAM (protospacer adjacent motif)-interacting domain (PID), a RuvC nuclease domain and an HNH nuclease domain (Figure 1A). Cas9 searches for and initially locates a PAM by its PID, which leads to unwinding of the protospacer and formation of the DNA-sgRNA heteroduplex, or R-loop. Finally, the HNH domain cleaves the target strand while the RuvC domain cleaves the non-target strand^22–24^. Previous biochemical and biophysical studies on SpyCas9 showed that it is essentially a single turnover enzyme with the rate-limiting step at DNA binding^25^ or R-loop formation^26^. Importantly, DNA cleavage efficiency scales with sampling of the HNH domain conformation; a correct conformation enhances the cleavage activity of the RuvC domain^27^. This allostery between HNH and RuvC is believed to result from the hinge comprising two helices that connect RuvC-III to HNH^21, 27–32^. Mutations that disrupt one of the two hinge helices lowered the DNA cleavage activity of SpyCas9^27^. These detailed mechanistic studies suggest that the allosteric hinge is also a potential target for engineering CECas9^33^. However, protein engineering by rational design to select for CECas9 variants would be laborious and have limited chances of successes.

**Figure 1:**
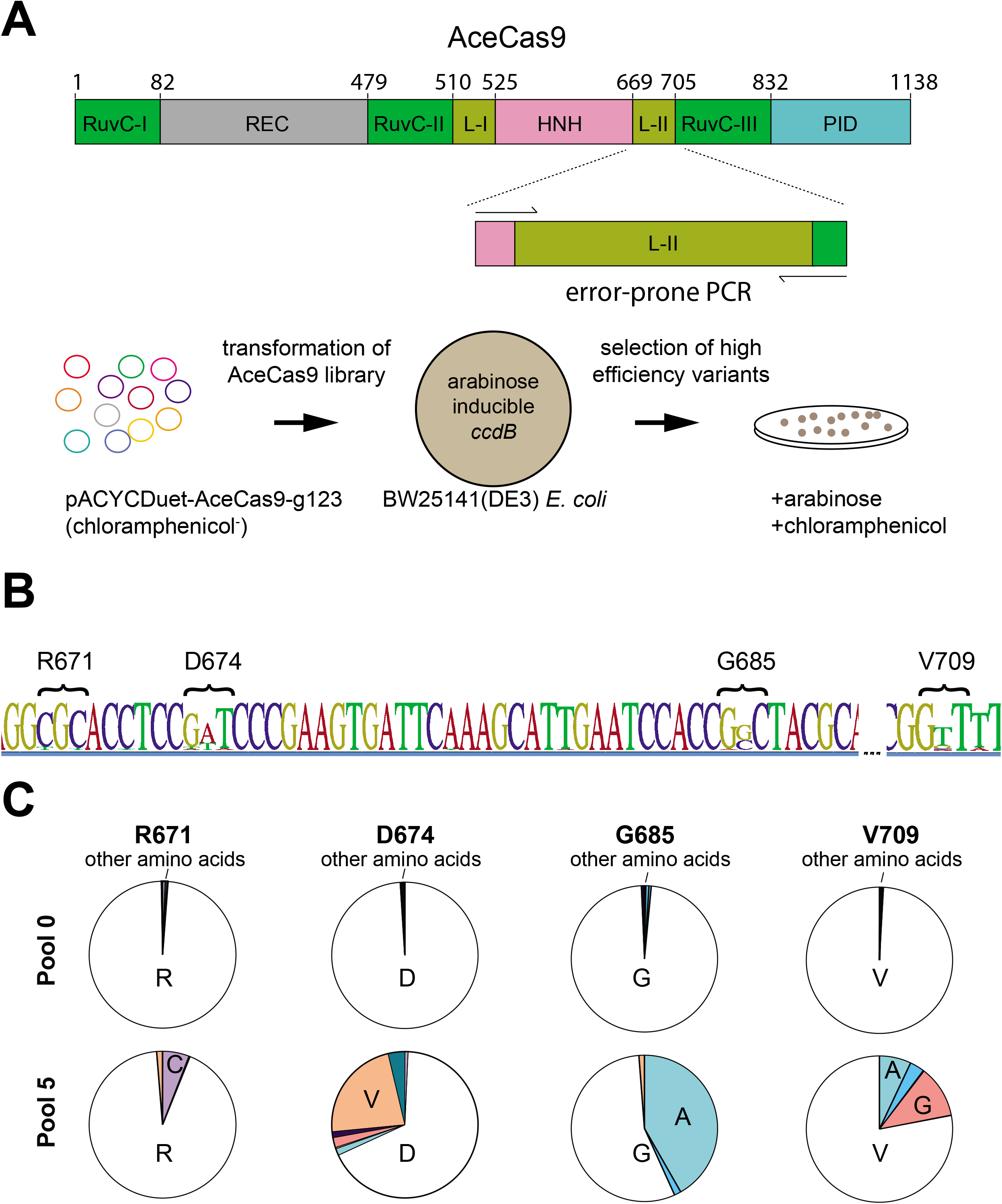
Selection of catalytically enhanced AceCas9. **A.** Top, Domain organization of AceCas9 with amplified view of linker II (L-II) with primers used for error-prone PCR primers. Bottom, Directed protein evolution selection strategy. pACAY-AceCas9-g123 and pACAY-AceCas9-g119 co-express AceCas9 and the single guide RNA with 24mer and 20mer guide, respectively. **B**. Sequence logo of consensus sequence from NGS analysis of final pool. The height of each letter is proportional to the observed frequency of each nucleotide in the alignment column. The locations with highest deviation are marked by their corresponding amino acid positions. **C.** Pie charts illustrate enrichment of amino acids at the four highest frequency sites. Pool 0 reflects enrichment values in the starting library whereas pool 5 reflects those from the last round of survival colonies. Single letter for amino acids are used.

We previously described a toxicity-based bacterial survival assay that is amenable for a library-based directed protein evolution selection^34, 35^. In this approach, a library of plasmids encoding a collection of Cas9 variants is transformed into bacterial cells harboring a plasmid expressing the toxic protein ccdB under the induction by arabinose. In presence of arabinose, cells transformed with the Cas9 variants capable of cleaving the ccdB-encoding plasmid would grow whereas those containing the variants incapable of cleaving the ccdB-encoding plasmid would not. Thus, the survival cell colonies would allow identification of the Cas9 variants that cleave the ccdB-encoding plasmid. To identify CECas9 variants, we need to program the ccdB-encoding plasmid with a protospacer that is not cleaved by the wild-type Cas9.

AceCas9 was previously shown to be most efficient in cleaving protospacers with 24 nucleotide (nt) of spacer complementarity^36^. The rate of single-turnover cleavage of the 24-nt protospacer is 4-fold that of the 20-nt protospacer on supercoiled plasmid DNA. In bacterial cells, while AceCas9 targeting a 24-nt protospacer causes near one hundred percent survival in the ccdB-based cell survival assay, that targeting a 20-nt protospacer has less than 0.2% rate of survival (Figure S1). In a typical transformation experiment, therefore, no surviving colonies would be observed when the wild-type AceCas9 is introduced to electro-competent *E. coli* cells harboring the ccdB-plasmid with a 20-nt protospacer^35, 36^. We took advantage of this selection feature in identifying AceCECas9. We hypothesized that the allosteric hinge can be engineered to allow a faster HNH conformational sampling, which leads to faster RuvC cleavage, resulting in more efficient formation of double-stranded breaks. To this end, we generated a library of the hinge variants through error-prone Polymerase Chain Reaction (PCR) (Figure 1A).

In contrast to negligible growth of cells containing the ccdB plasmid transformed with a 20-nt protospacer wild-type AceCas9, those transformed with the hinge library showed significant growth. The survival colonies were then pooled, and their plasmid DNA were extracted for a second round of selection. The selection cycle was repeated until the rate of survival per microgram of DNA no longer increased. The DNA pool from each selection cycle was subjected to Next Generation Sequencing (NGS) and plasmid DNA from 20 individual colonies of the final cycle were also isolated for Sanger sequencing. We identified the survival variants via Single Nucleotide Polymorphism (SNP) calling on the assembled contigs of each DNA pool (Figure 1B & 1C). The most deviations occur at the nucleotide positions corresponding to amino acid Arg^671^, Asp^674^, Gly^685^, and Val^709^(Figure 1B & 1C). These survival variants encode AceCECas9 R671C, D674V, G685A, and V709A/G. The findings were supported by Sanger sequencing of individual colonies, which showed 11/20 to be G685A, 6/20 to be V709A/G, and 3/20 to be D674V. We mapped these amino acid positions onto our previously constructed AceCas9 structure model (Figure S2). Arg^671^, Asp^674^, and Gly^685^ are located downstream of the helix following the RuvC domain (helix 1) (AceCas9, like other type II-C Cas9s, lacks the second helix following helix 1). Surprisingly, V709 is located outside the predicted hinge region on the ß-strand directly packing against hinge helix 1 (Figure S2), suggesting that structural elements interacting with the hinge helices also impact the allosteric regulation.

In order to confirm the functional activities of the selected AceCECas9, we made individual AceCas9 mutants of G685A and V709A and tested both in-cell and in vitro cleavage assays. Not surprisingly, G685A and V709A had nearly 100% survival rate in *E. coli* cells harboring the ccdB-plasmid with the 20-nt protospacer (Figure 2A) while the wild-type AceCas9 would not survive (Figures 2 & S1). Consistently, the single-turnover rate of cleavage for both mutants are 3-4-fold of that of the wild-type AceCas9 for either the 24-nt (24mer) or the 20-nt (20mer) protospacer (Figure 2B). Thus, the significantly increased rate of cleavage allows AceCECas9 to effectively eliminate the toxic ccdB-plasmid with a 20-nt protospacer, promoting cell survival.

**Figure 2:**
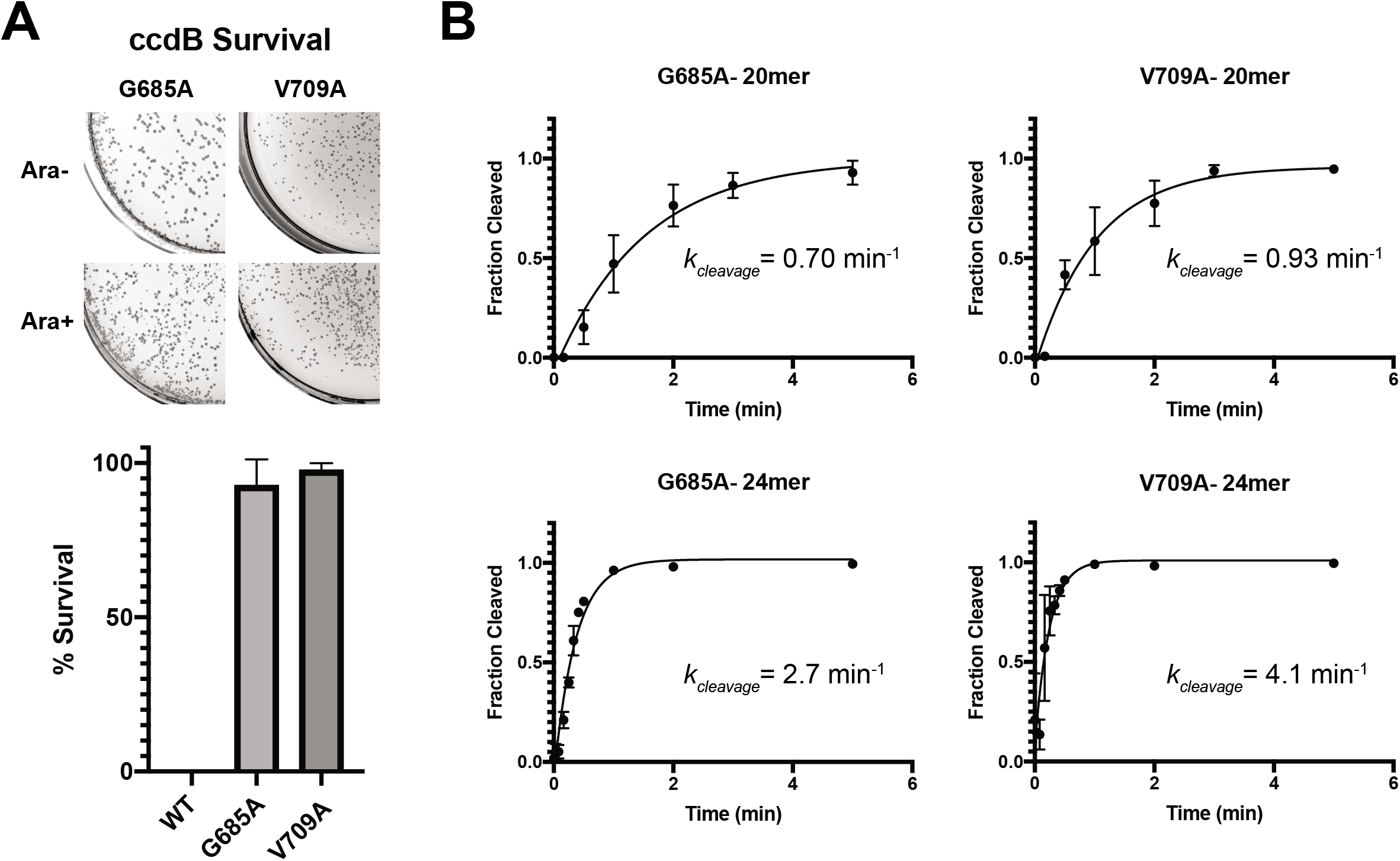
Verification of the catalytically enhanced AceCas9 by survival assay and kinetic analysis. **A,** Plasmids encoding the selected AceCas9 mutants were transformed into ccdB harboring cells that are plated on plates with (Ara+) or without arabinose (Ara-) respectively. Images of the plates are shown with the corresponding survival rate plotted below. **B,** Single-turnover DNA-cleavage experiments and the resulting rates of the AceCECas9 with either the 20mer spacer or the 24mer spacer sgRNA. Molar excess of AceCas9-sgRNA complex were incubated with plasmid DNA for various times at 50 °C before being resolved and visualized on agarose gels. Kinetic experiments were performed in triplicate.

In order to evaluate if AceCECas9 variants have increased tolerance of DNA-sgRNA mismatches, we assessed the survival rates of G685A and V709A in *E. coli* cells harboring the ccdB-plasmids with mismatched protospacers at position −1 or −4. These mismatches were previously shown to prevent the wild-type AceCas9 from eliminating the ccdB-plasmid thus leading to death of the *E. coli* hosts^35^. Similarly, neither G685A nor V709A survived the T(−1)G plasmid (Figure S3), although a small percentage of cells survived for the G(−4)A protospacer, suggesting that AceCECas9 does not significantly escape seed region mismatches in comparison with the wild-type.

In order to identify the molecular basis for the enhanced catalytic efficiency, we performed DNA binding competition experiments to learn if AceCECas9 enhances DNA binding, DNA unwinding, or HNH conformational sampling. To a typical DNA cleavage assay containing 6 nM plasmid target and 250 nM AceCas9 (or AceCECas9), doublestranded DNA oligo competitors were added at increasing concentrations. The fraction of cleavage was then quantified and plotted against the logarithm of competitor concentrations. The plot was then fitted using a standard competition binding model to yield an apparent competitor binding constant, K_I_. The wild-type protospacer DNA oligo has nearly identical K_I_ for both AceCas9 and AceCECas9 (Figure 3A). The T(−1)G protospacer DNA oligo is a worse competitor than the wild-type but again with equal K_I_ for both AceCas9 and AceCECas9, suggesting that AceCECas9 does not enhance, in comparison with AceCas9, DNA binding for either the wild-type or mismatched protospacer (Figure 3A). Finally, AceCas9 and AceCECas9 showed similar binding constants to two bubbled DNA oligos (either complementary or not to sgRNA) (Figure 3B), indicating that the DNA unwinding was not altered in AceCECas9. These results confirm that the enhanced catalytic efficiency of AceCECas9 is through a process other than DNA binding and unwinding, most likely HNH domain sampling.

**Figure 3:**
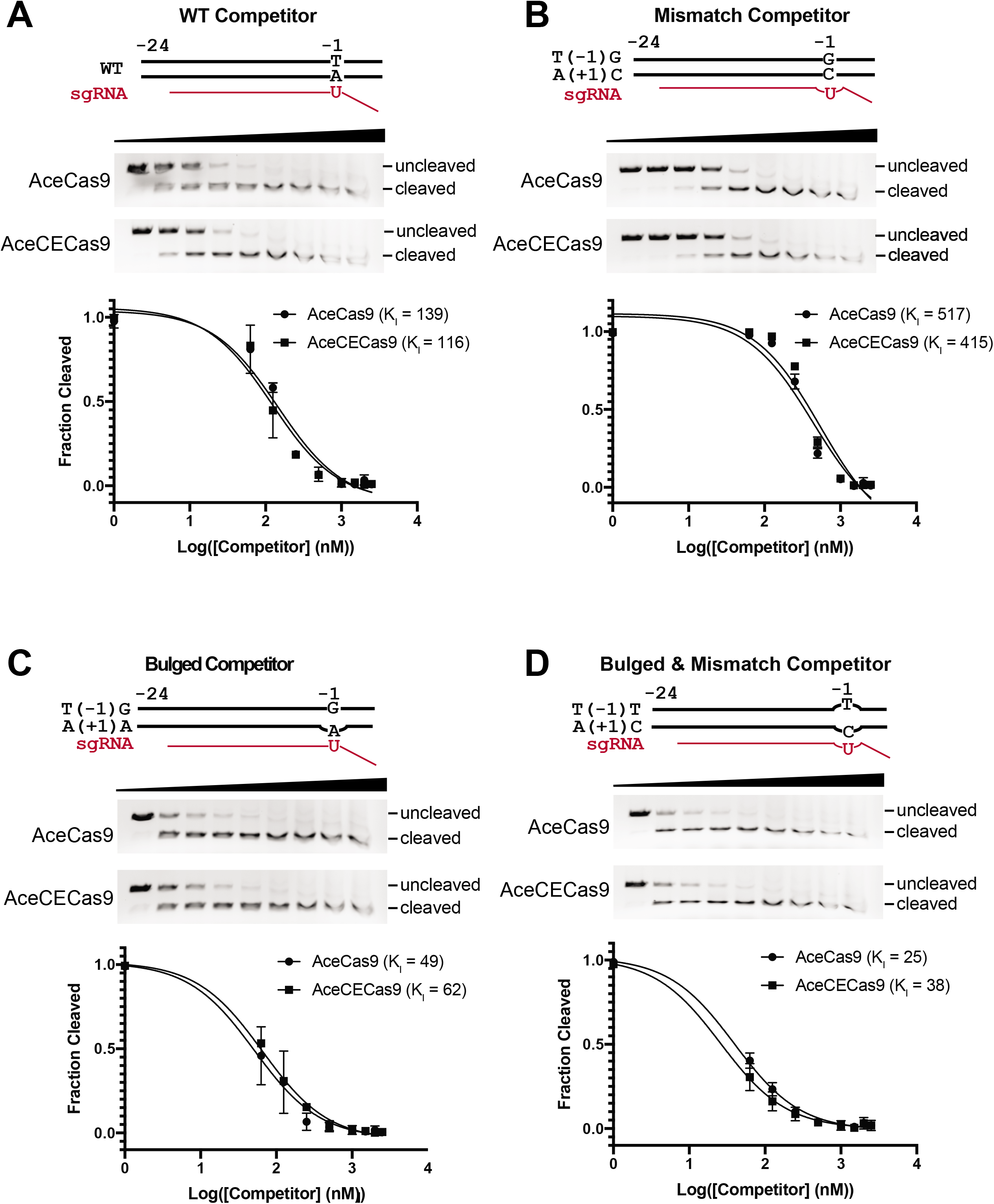
DNA binding competition assay results. The DNA oligo substrates used are schematically shown as wild-type (WT), T(−1)G/A(+1)C (dsDNA with mismatch to sgRNA at position +1), T(−1)T/A(+1)C (bulged DNA oligo at position +1 that mismatches with sgRNA), and non-target strand bulge (bulged DNA oligo at position at+1 that matches sgRNA). For each competition binding experiment, the cleavage gel image and fraction of cleavage versus competitor concentration at logarithmic scale are shown. Solid curves are fitted theoretical curves and fitted K_I_ values are indicated. **A,** Competition binding result for the wild-type or T(−1)G oligo with AceCas9 (WT) and the V709A variant. **B,** Competition binding result for the bulged oligos with AceCas9 (WT) and the V709A variant.

Finally, we applied the same directed evolution strategy to the more widely used SpyCas9. Similar to AceCas9, we systematically reduced the base pairs in the protospacer complementary to the guide RNA and determined that 17 base pairs slowed the wildtype SpyCas9 sufficiently to obstruct survival. We then generated a DNA library that encodes SpyCas9 hinge variants (with guide RNA) and transformed it to the cells harboring the *ccdB* plasmid with a 17-base pair protospacer. Unlike the wild-type SpyCas9 transformation that resulted in no cell survival, the library transformation resulted in significant growth on arabinose containing plates, and after iterative rounds of evolution, a single variant was identified by sequencing (Figure S6). We termed the variant catalytically enhanced SpyCas9 (SpyCECas9). To determine if the SpyCECas9 improves gene editing applications relative to the SpyCas9, we employed both to achieve homology directed repair (HDR)-based insertion of the green fluorescence protein (GFP) gene in the LMNB1 gene within HCT116 cells (Figure 4 & Figure S7). As shown in Figures 4C, 4D & 4E, co-transfection of the SpyCECas9-encoding plasmid with the HDR template into HTC116 consistently resulted in 2-fold increase in GFP incorporation versus that of SpyCas9, likely due to its enhanced cleavage of the target DNA.

**Figure 4:**
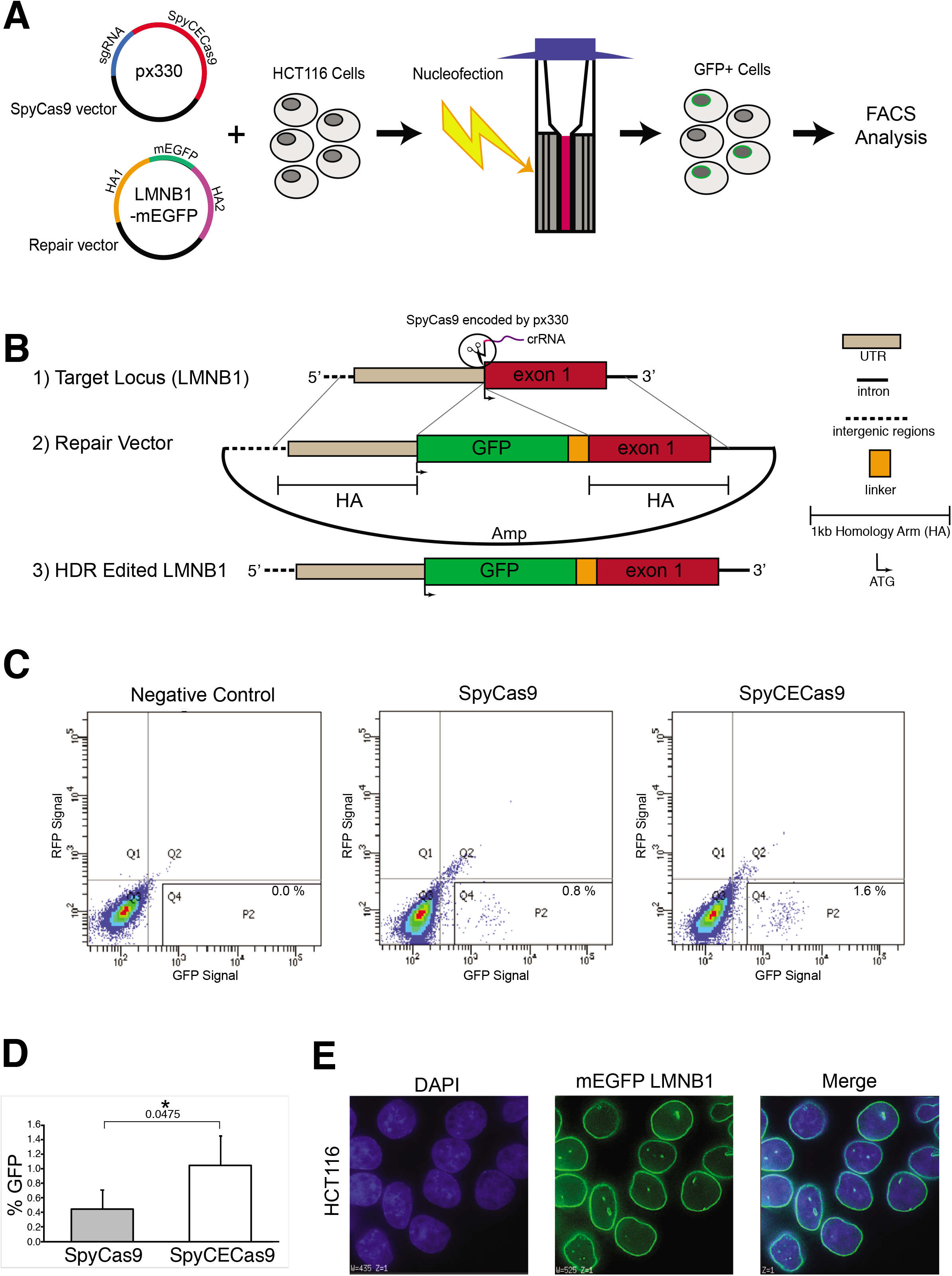
Gene insertion facilitated by the wild-type and the catalytically enhanced SpyCas9 in HCT116 cells. **A,** Cartoon illustration depicting the genome insertion process. The px330 vector co-expressing either the wild-type SpyCas9 or the SpyCECas9 with the guide sgRNA targeting the LMNB1 gene is co-transfected with the repair vector containing mEGFP flanked by 1kb homology arms (HAs). Each transfection is performed with the Lonza SE cell line 4-D Nucleofector solution kit by electroporating one million HCT116 cells in an electro-cuvette. Cells are plated and cultured for 48-hours post transfection (positive cells are depicted as grey with a green ring around the nuclear lamina). The HCT116 cells that successfully achieved homology directed repair (HDR) by incorporation of the green fluorescent protein (GFP) are detected, sorted, collected via flow cytometry and expanded for imaging. **B,** Detailed scheme of the CRISPR-Cas9-mediated green fluorescence protein (GFP) tagging strategy. 1) The crRNA (purple) co-expressing with the wild-type SpyCas9 (px330, Addgene #42230) or the catalytically enhanced (CE) SpyCECas9 on a modified px330 vector targets a site upstream of exon 1 (red) of the LMNB1 gene. 2) A repair vector containing 1kb homology arms (HAs) flanking the GFP-encoding sequence fused with a linker sequence (AICSDP-10:LMNB1-mEGFP, Addgene #87422). 3) Successfully edited target locus following SpyCas9 cleavage and homology directed repair contains an in-frame insertion of the GFP after the start codon of LMNB1 exon 1. **C,** The observed GFP signal of the wild-type and the catalytically enhanced SpyCas9. Flow cytometry plots displaying GFP fluorescence intensity (x-axis) versus RFP fluorescence intensity (y-axis; to control for auto-fluorscence) following 48-hour posttransfection for the wild-type (SpyCas9) and catalytically enhanced (SpyCECas9). GFP positive cells fall within the P2 gate. The negative control is obtained from cells transfected with the repair vector and SpyCas9 without a sgRNA. Experiments were conducted in triplicate (Figure S7). **D,** Bar graph displaying the averages of the quadruplicate measurement for both SpyCas9 and SpyCECas9 and error bars displaying standard deviation. Statistical significance was determined by an unpaired t-test (\**p*≤0.05, exact value shown below the asterisk). **E,** Representative fluorescence microscopy images of GFP-tagged LMNB1 in HCT116 cells obtained on a DeltaVision (GE Life Sciences) microscope. The cell nuclei were stained with DAPI.

Our results present the first engineered Cas9s with increased catalytic efficiency through a directed protein evolution method. We demonstrated the effectiveness of this method on AceCas9 that yielded enhanced DNA cleavage in bacterial cells. We then used the same strategy to produce SpyCECas9 that doubled the efficiency of the SpyCas9-mediated HDR in human colon cancer cells. The catalytically enhanced Cas9s can be applied in areas where they are currently ineffective.

## Methods

### Plasmid and Library Construction

The arabinose-inducible *ccdB-encoding* plasmid (p11-LacY-wtx1) was a gift from Dr. David Edgell (University of Western Ontario). The target sequence was inserted downstream of the *ccdB* gene using restriction enzymes. Point mutations were made via Q5 site-directed mutagenesis (NEB).

Genes encoding AceCas9 and the sgRNA were inserted into the pACYCDuet vector (Novagen 71147) to form pACYCDuet-Ac9-g119 (20mer) or pACYCDuet-Ac9-g123 (24mer), respectively, as previously described^36^. The hinge-library was created by amplifying the predicted hinge region with Gibson assembly primers via error prone PCR (Ser^663^-Gly^712^) at an average rate of 1 mutation per 100 bp and inserting the PCR products into the pACYCDuet-Ac9-g119 vector. The resulting DNA was transformed into DH5α cells. About 60,000 colonies were obtained after overnight growth on media containing chloramphenicol. The colonies were pooled, grown for 2 hours, then collected for extraction of plasmid DNA.

### In vivo selection

The survival assay was carried out as previously described^34–36^. Briefly, electrocompetent *E. coli* BW25141 containing the modified p11-LacY-wtx1 (ccdB encoding) plasmid were transformed with 100-200 ng of AceCas9 library plasmids. Cells were then recovered in prewarmed Super Optimal Broth (SOB) for 30 minutes with shaking at 37 °C before 0.05 mM IPTG was added, and recovery continued for an additional 60 minutes. Transformants were then plated on LB agar plates containing either chloramphenicol or chloramphenicol and 10 mM arabinose and incubated at 37 °C for 16-20 hours. Colonies were counted manually, and survival rates were calculated by dividing the colony forming units (CFU) of arabinose-containing plates by that of the chloramphenicol only plates.

### Next Generation Sequencing

The hinge region of each survival pool was PCR amplified with primers containing Illumina indexes. Adaptors were added to these PCR products and then the pooled libraries were subjected to 300 cycle single-end sequencing using MiSEQ (Illumina). Data was analyzed using Geneious 9.0.5 NGS analysis tools.

### Protein Expression and Purification

AceCas9 and mutant variants were expressed and purified as previously described^36^. Briefly, harvested cells in a lysis buffer were lysed via sonication. Clarified lysate was loaded onto an Ni-NTA agarose (Qiagen) column. Elutant was further purified via a HiTrap Heparin-HP (GE Healthcare) column and then a size exclusion column (HiLoad 26/60 Superdex 200 increase GE Healthcare) before concentration and storage at −80 °C.

### In-vitro transcription and Purification of sgRNA

DNA oligos coding for a T7 promotor followed by the sgRNA were annealed in 10x PCR buffer by heating in boiling water for 5 minutes followed by slow cooling. In-vitro transcription was performed by adding 1 uM of this dsDNA to 50 mM Tris pH 8.0, 40 mM MgCl_2_, 10 mM dithiothreitol (DTT), 2 mM Spermindine, 5 mM ATP, UTP, CTP, and GTP pH 8.0, 0.1% Triton X-100, and 0.15 mg/mL of homemade T7 RNA Polymerase, and incubating at 37 °C overnight. The reaction was quenched the following morning by adding 50 mM EDTA. The sgRNA was then extracted via phenol/chloroform pH 4.5 extraction and purified via 10% denaturing PAGE.

### Plasmid Cleavage Assay

Plasmid cleavage assays were performed as previously described^34–36^. Briefly, 500 nM pre-annealed AceCas9: sgRNA complex was incubated with 6 nM plasmid DNA in cleavage buffer (20 mM Tris pH 7.5, 150 mM KCl, 2 mM DTT, 10 mM MgCl_2_, 5% glycerol) at 50 °C for a given time. Reactions were then stopped by adding 5x stop buffer (25 mM Tris pH 7.5, 250 mM EDTA, 1% SDS, 0.05% w/v bromophenol blue, 30% glycerol) and resolved on a 1x TBE 0.5% agarose gel containing Ethidium bromide. Gels were imaged by the ChemiDoc XRS System (Bio-Rad).

### Single Turnover Kinetics

Plasmid kinetics assays were performed in a similar manner as the plasmid cleavage assays, but all reagents (DNA, RNP complex, and Eppendorf tubes) were kept ice cold before reactions were initiated. After initiation of reaction, samples were taken from 50 °C water bath in triplicate at each time point and placed on ice, then ice cold stop buffer was quickly added. Gels were analyzed by Image Lab Version 5.2.1 build 11 (Bio-Rad) and fitted to an exponential plateau curve in Prism 8.1.1 to extract k_cleavage_.

### DNA Binding Competition Assays

Oligonucleotides were annealed by heating in 95 °C water bath followed by slow cooling. Serial dilutions of oligos were made from 25 uM to 0.063 uM, and added to 6 nM plasmid DNA. Plasmid/oligo mixture was then added to annealed RNP complex (250 nM AceCas9 (or AceCECas9) and 250 nM sgRNA) and allowed to react for 15 minutes at 50 °C. Reactions were stopped with stop buffer, resolved on agarose gels before being visualized, and quantified as before.

### CRISPR-mediated Homology Directed Repair (HDR) Assay in HCT116 cells

HCT116 cells were maintained in McCoy’s 5A media supplemented with 10 % fetal bovine serum and 1 % penicillin/streptomycin at 37 °C with 5% CO2. The plasmids encoding the wild-type SpyCas9 (px330, Addgene #42230) or its catalytically enhanced variant, SpyCECas9 (modified px330), targeting the LMNB1 gene (GGGGTCGCAGTCGCCATGGC) (2 ug) and an LMNB1-mEGFP containing repair vector (Addgene plasmid #87422) (3 μg) were nucelofected into HCT116 cells grown to 70-80% confluency using the Lonza SE cell line 4-D Nucleofector solution kit with pulse code EN113. 48-hours post-nucleofection, cells were evaluated for the presence of GFP using FACSCanto.

## Acknowledgments

We thank D. Edgell for providing cells and plasmids for in vivo study; B. Washburn, C. Pye, and Kristina Poduch of the FSU Molecular Cloning Facility for the cloning experiments and discussion; S. Miller and A. Brown of the FSU Sequencing facility for assistance with NGS library preparation and sequencing. This work was supported by NIH grants R01 GM099604 to H.L. and R01 GM083337 to D.M.G.

## Author Contributions

T.H. and H.L. designed all experiments. T.H. designed primers for *ccdB* plasmid variants and AceCas9 plasmid libraries, created AceCas9 plasmid libraries, prepared DNA for NGS and analyzed results, purified AceCas9 and variants obtained from library screens, and performed in vitro DNA cleavage assays. M.R., C.S., and E.S. performed in vivo survival assays with AceCas9 plasmid libraries for iterative rounds, prepped DNA for each pool, and analyzed Sanger sequencing results. M.R. and K.K. transfected HCT116 cells and analyzed results. E.S. designed primers for variants obtained from library screens. T.H. and H.L analyzed data, wrote and edited manuscript and figures.

---

**Supplemental Figure 1:**
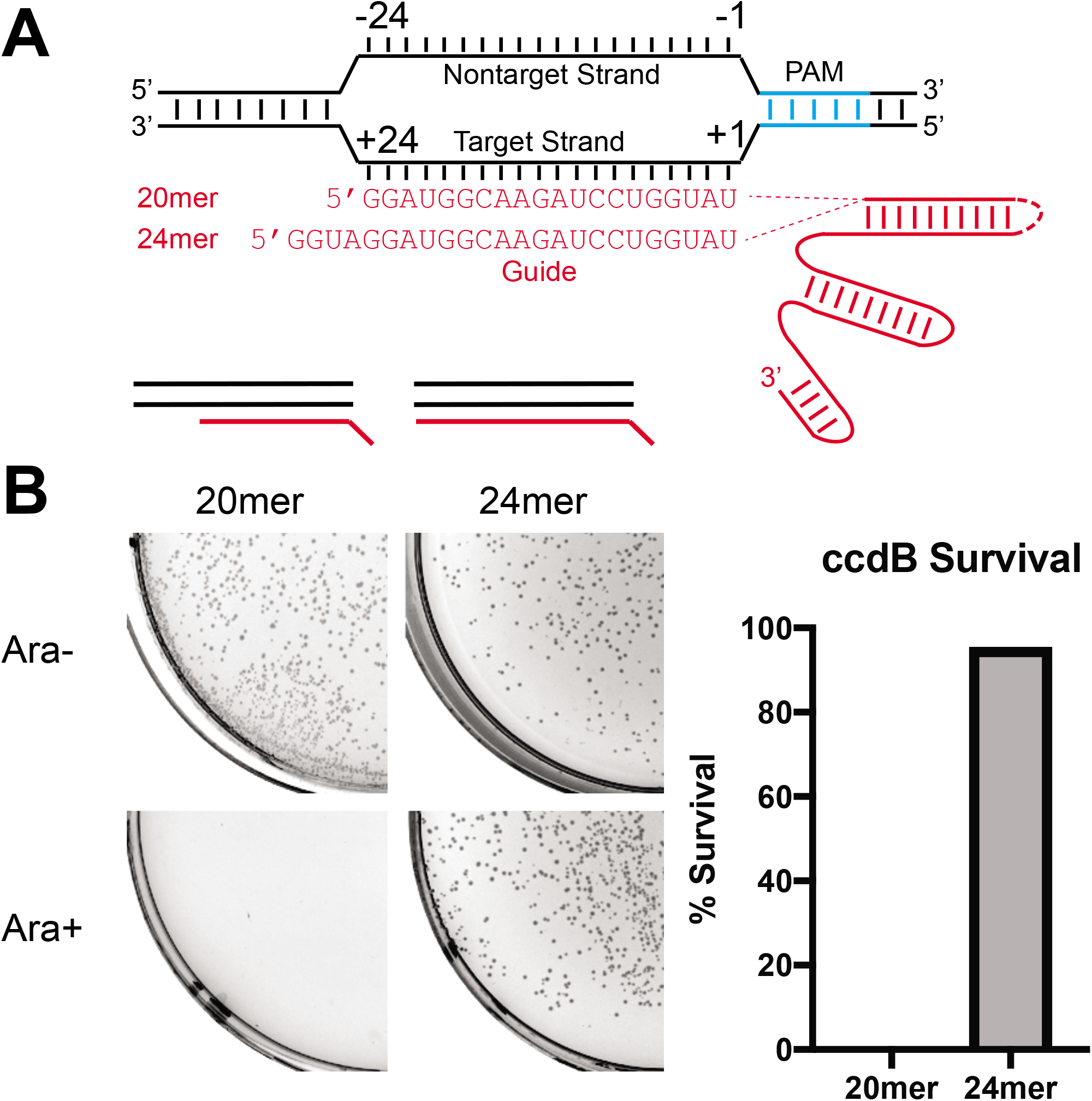
Sensitivity of AceCas9 to spacer length. **A,** Structural features of target DNA and sgRNA used in this study. The numbering of the terminal protospacer nucleotides are indicated. **B,** Plasmids encoding AceCas9 with either 20mer (pACAY-AceCas9-g119) or 24mer (pACAY-AceCas9-g123) guide RNA were transformed into ccdB harboring cells. Images of plates in the absence (Ara-) and the presence of arobinose (Ara+) are shown with caculated survival rates plotted as bar diagrams to the right.

**Supplemental Figure 2:**
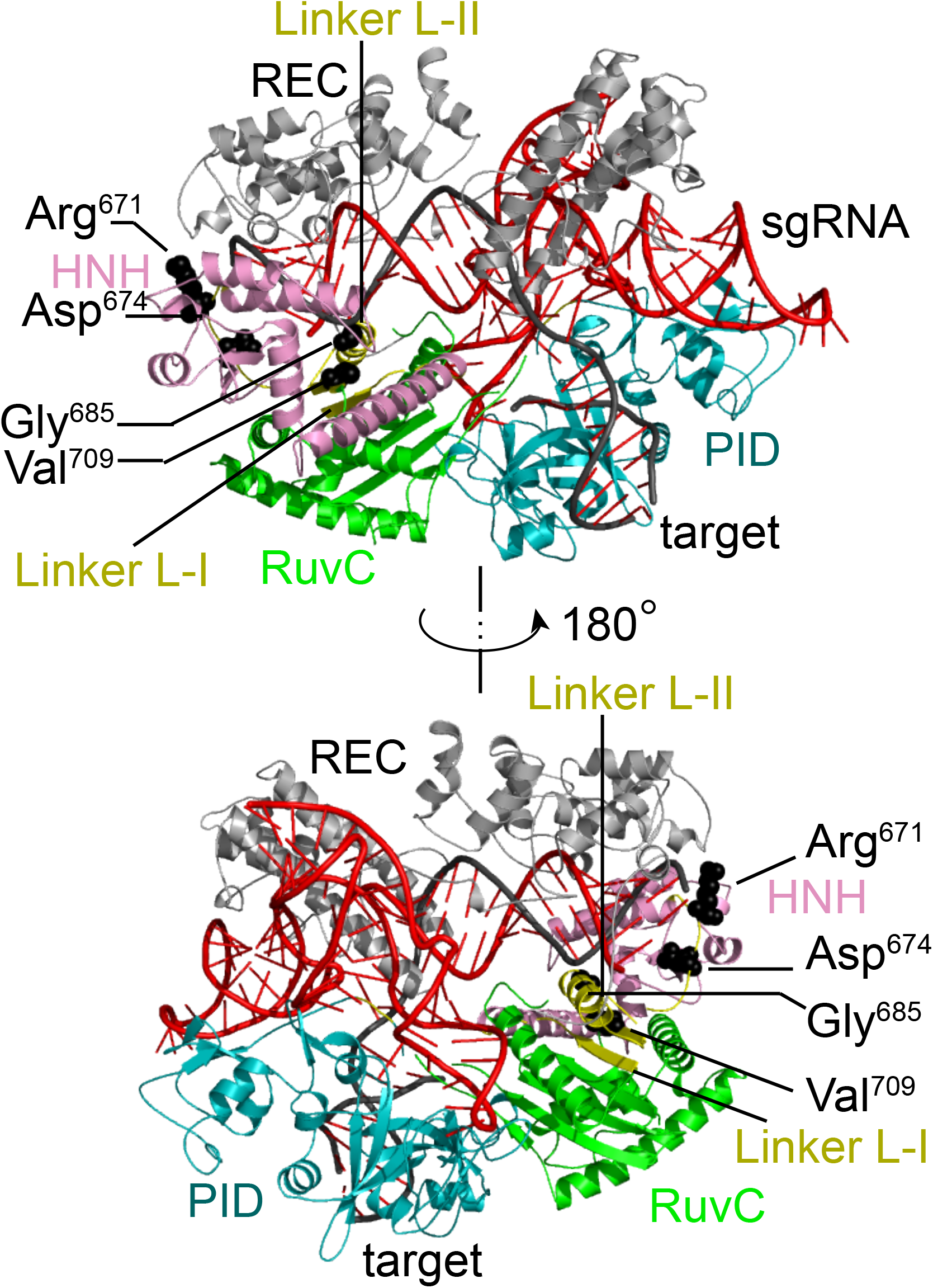
Mapping catalytically enhanced mutations on a modeled AceCas9 structure. AceCas9 domains are colored and labeled. Target DNA is shown in black and sgRNA shown in red. The mutations are shown as black spheres as labeled. REC, nucleic acid recognition domain (gray); HNH, HNH catalytic domain (pink); RuvC, RuvC catalytic domain (green); PID, PAM interacting domain (teal), and Linkers L-I and L-II (yellow).

**Supplemental figure 3:**
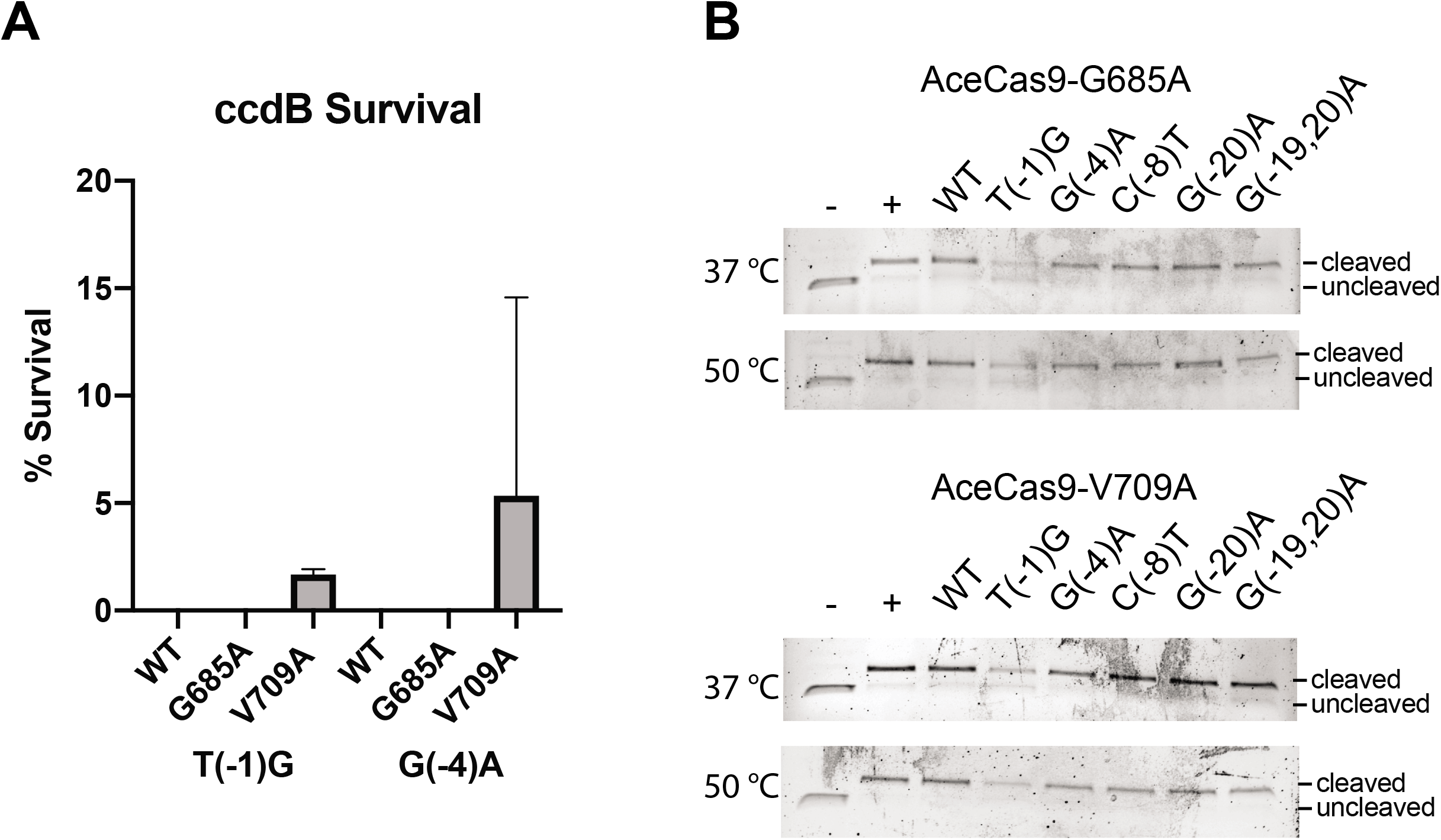
Off-target analysis of AceCECas9. **A,** Plasmids encoding AceCas9 (WT or AceCECas9 variants) were transformed into ccdB T(−1)G or G(−4)A harboring cells. Caculated survival rates shown in a bar graph. Error bars represent standard deviation (S.D.) from triplicates. **B,** in vitro cleavage reactions with AceCECas9 variants with plasmid DNA harboring mismatches in the protospacer. Molar excess of AceCECas9 was incubated with plasmids for 60 minutes at 50 °C or 37 °C before being separated and visualized on agarose gels.

**Supplemental Figure 4:**
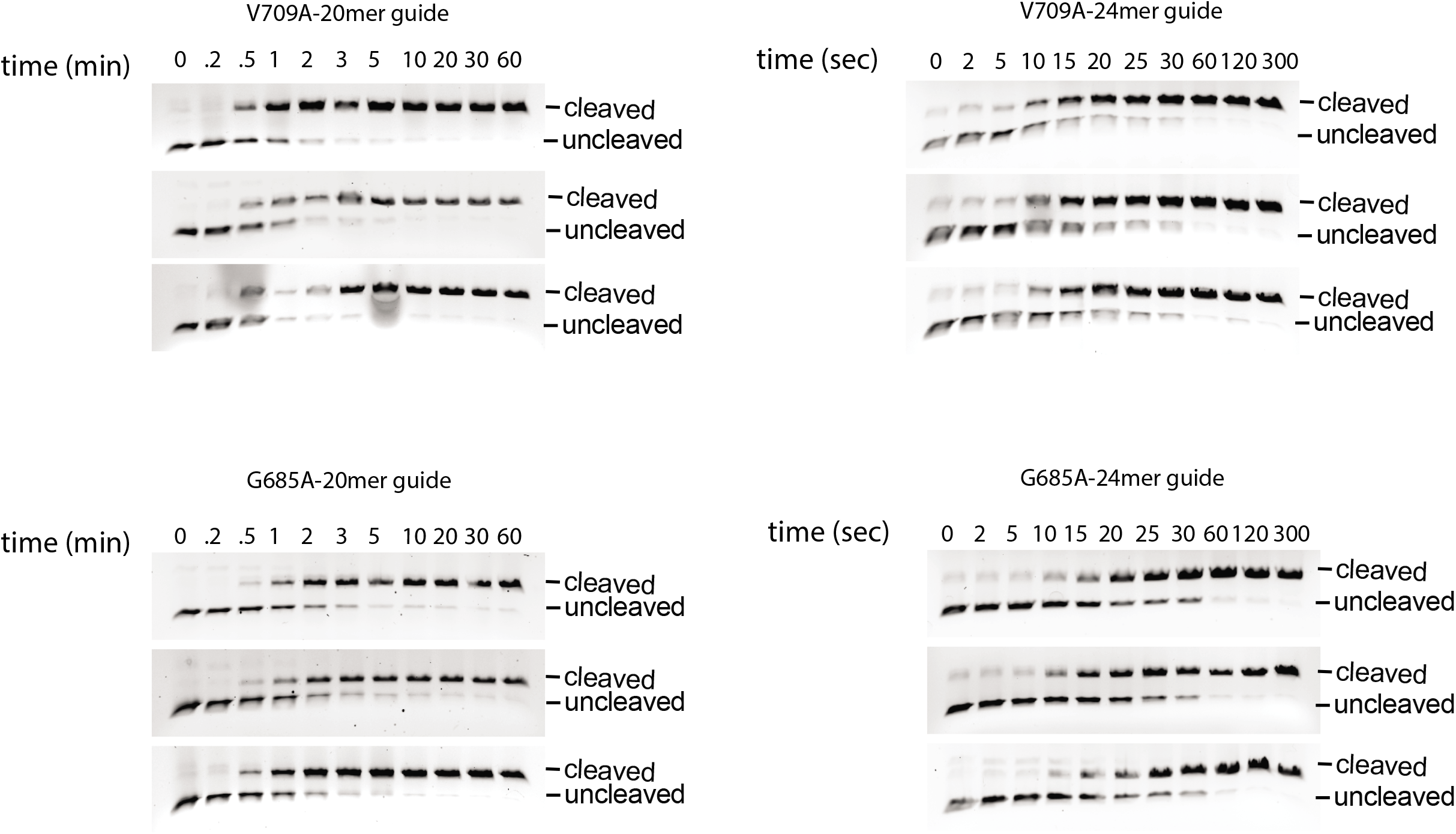
Single turnover cleavage assay gel images that were used to produce binding curves in Figure 2. Molar excess of AceCECas9 with either 20mer guide or 24mer guide RNA were incubated with plasmid DNA for increasing time points. The cleavage products were resolved and visualized on agarose gels.

**Supplemental Figure 5:**
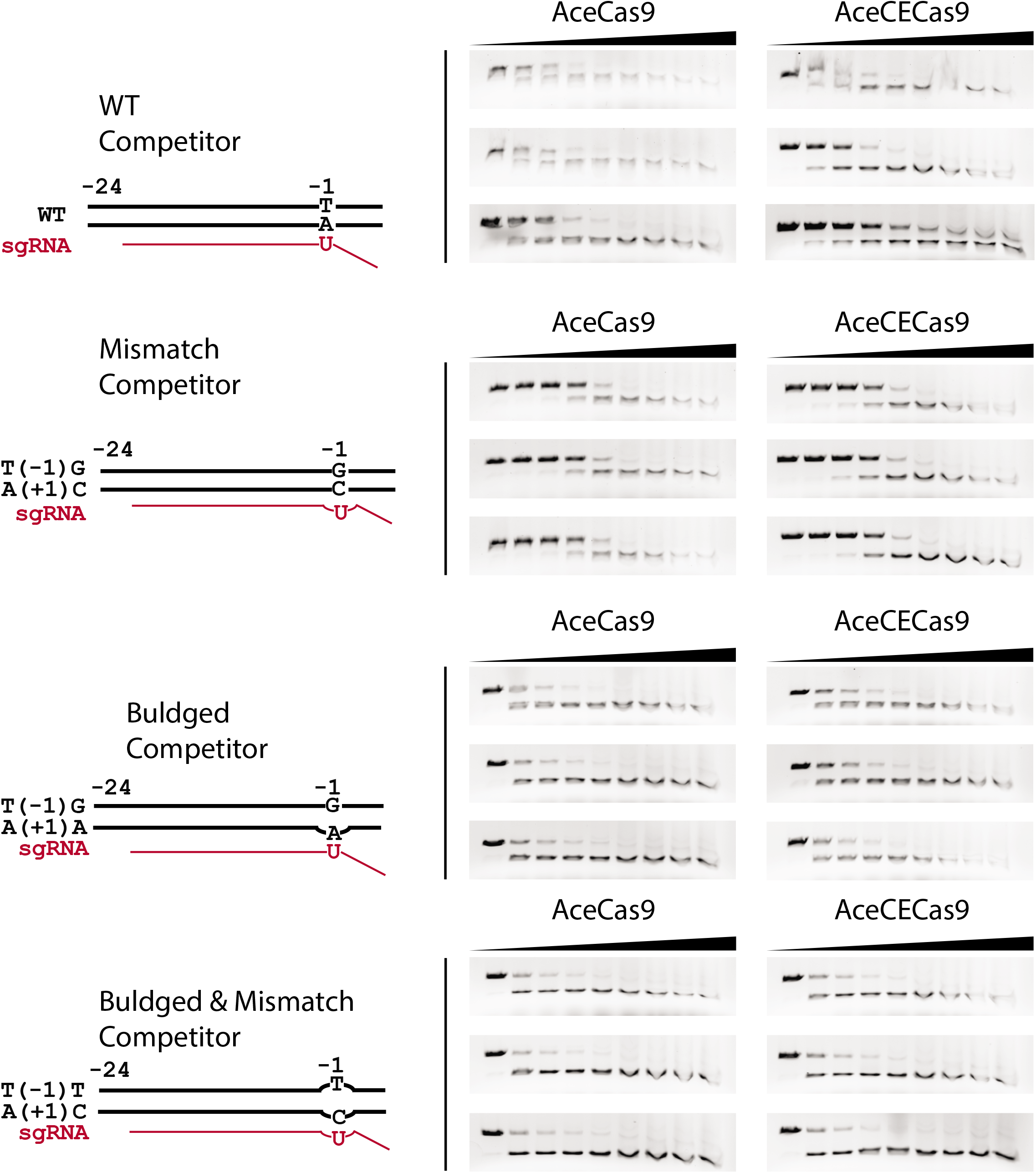
Compeition binding gel images that were used to produce the binding curves in Figure 3. Molar excess of the catalytically enhanced (CE) AceCas9, V709A, with the 24mer guide RNA incubated with plasmid substrate DNA were in the presence of increasing concentrations of dsDNA oligo competitors (0 – 2.5 μM) for 15 minutes at 50 °C. Triplicates of the cleavage proucts were resolved, visualized on agarose gels and averaged for curve fitting.

**Figure S6.**
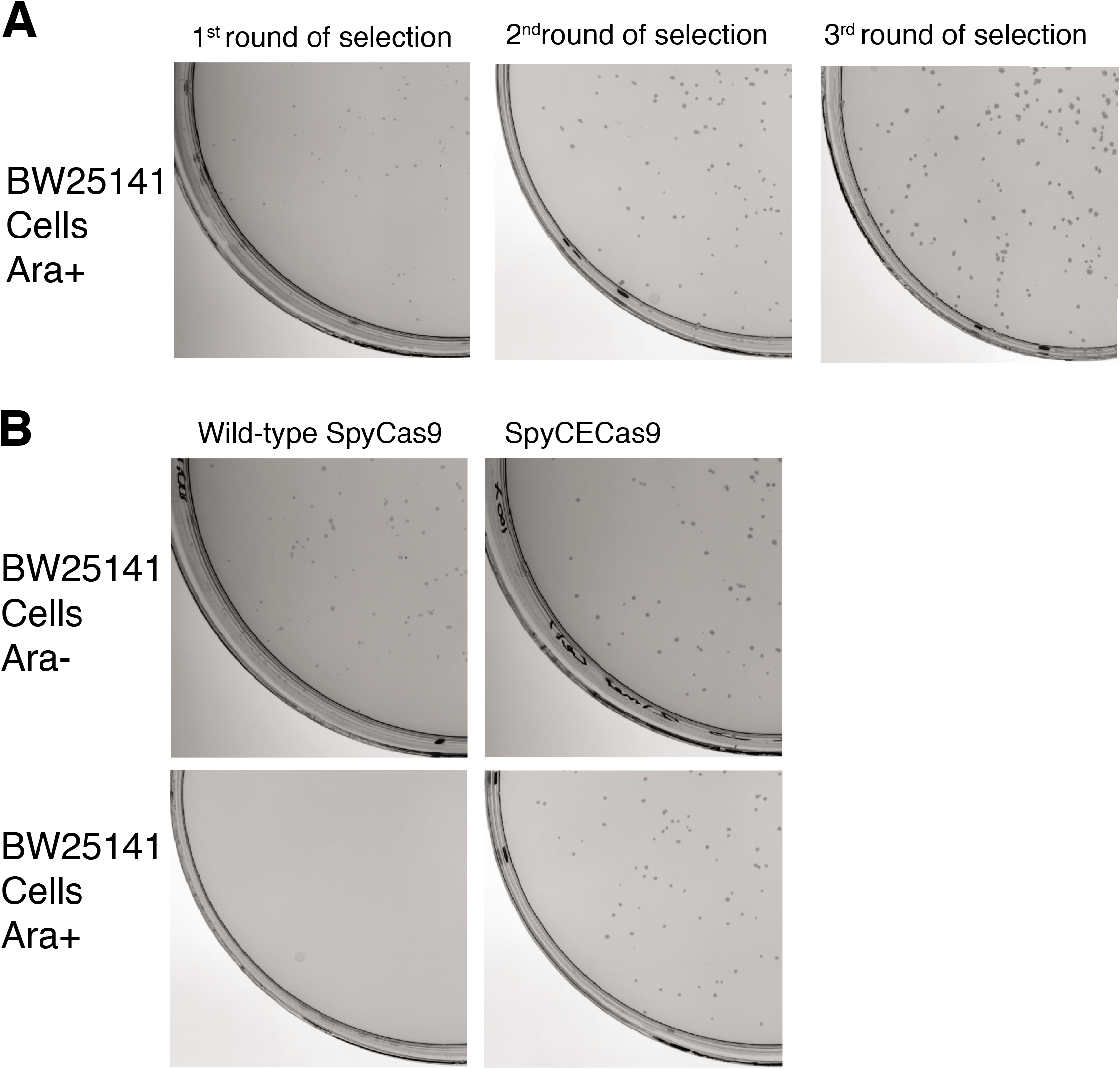
Directed protein evoution of catalytically enhanced SpyCas9 (SpyCECas9). **A.** 0.1 μg DNA plasmid encoding a library of SpyCas9 containing variants at its hinge helices were transformed into BW25141 cells harbouring ccdB-encoding plasmids that bares a 17mer protospacer for SpyCas9. The entire recovery volume was plated on arbinose-con-taining (Ara+) plates. Images show the plates for the three rounds of selections. **B.** Confirmation of the selected SpyCECas9 variants by the cell toxicity assay. The wild-type or the plamsid encoding the catalytically enhanced SpyCas9 was transformed into BW25141 cells harboring the ccdB-encoding plasmid that bears a 17mer protospacer for SpyCas9. Plates contain either no (Ara-) or arabinose (Ara+).

**Figure S7.**
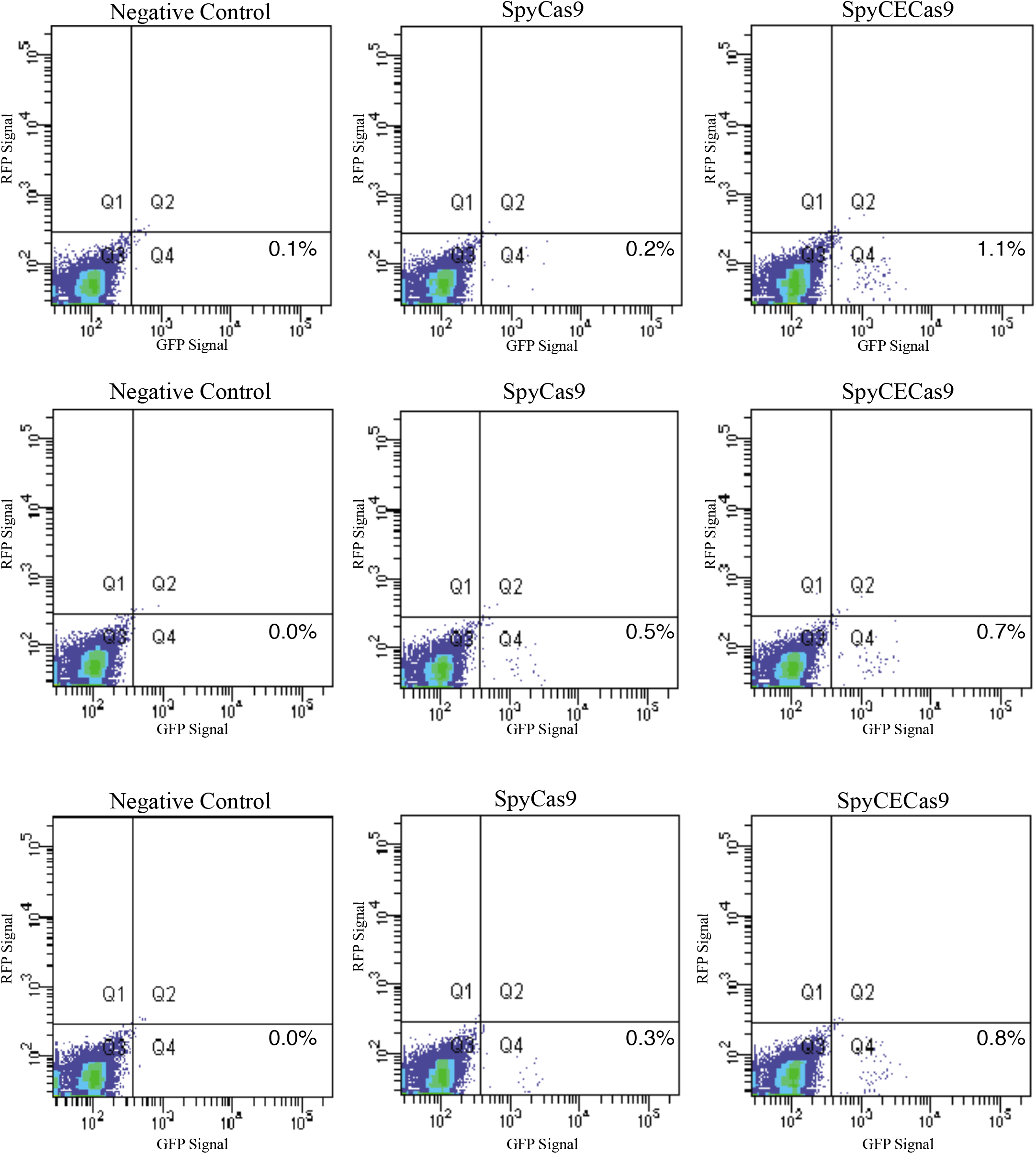
Flow cytometry counts for cells expressing green fluorescent protein (GFP). **A.** Triplicate experiments of SpyCas9-mediated homology directed repair (HDR) to incorporate GFP-encoding sequence into the LMNB1 gene. Flow cytometry plots displaying GFP fluorecence in cells (x-axis, Q4) following 48-hour post-transfection of the wild-type (Spy-Cas9) or the catalytically enhanced SpyCas9 (SpyCECas9). The negative control is obtained from cells transfected with the plasmid encoding SpyCas9 (without sgRNA) only.

